# Genetically Targeted Ratiometric and Activated pH Indicator Complexes (TRApHIC) for Receptor Trafficking

**DOI:** 10.1101/180141

**Authors:** Lydia A. Perkins, Qi Yan, Brigitte F. Schmidt, Dmytro Kolodieznyi, Saumya Saurabh, Mads Breum Larsen, Simon C. Watkins, Laura Kremer, Marcel P. Bruchez

## Abstract

Fluorescent protein based pH sensors are a useful tool for measuring protein trafficking through pH changes associated with endo-and exocytosis. However, commonly used pH sensing probes are ubiquitously expressed with their protein of interest throughout the cell, hindering the ability to focus on specific trafficking pools of proteins. We developed a family of excitation-ratiometric, activatable pH responsive tandem dyes, consisting of a pH sensitive Cy3 donor linked to a fluorogenic malachite green acceptor. These cell-excluded dyes are targeted and activated upon binding to a genetically expressed fluorogen activating protein, and are suitable for selective labeling of surface proteins for analysis of endocytosis and recycling in live cells using both confocal and superresolution microscopy. Quantitative profiling of the endocytosis and recycling of tagged β2-adrenergic receptor (B2AR) at a single vesicle level revealed differences among B2AR agonists, consistent with more detailed pharmacological profiling.

---

G-protein coupled receptors (GPCRs) comprise the largest family of membrane signaling receptors.^1^ These cell surface receptors play an important role in controlling homeostasis and cellular phenotypes by relaying extracellular stimuli to intracellular downstream targets. Defects in GPCR signaling and trafficking are associated with many human diseases, such as drug addiction and heart disease.^2,3^ This class of receptors is the dominant class among all drug targets.^4^ The search for new GPCR drug candidates has demonstrated that different agonists can exhibit bias towards different cell signaling pathways.^5^ The pharmaceutical significance of ligand bias has altered the methods of therapeutic screens and drug candidate evaluation.^6^ GPCR desensitization through endocytosis and its intracellular sorting, recycling, or degradation, are important mechanistic features that are associated with GPCR signaling.^7,8^ B2AR has been extensively studied, and is a classic test-bed for developing new procedures for drug classification, including the influence of drugs on B2AR’s endocytic itinerary. The measurement of trafficking of GPCRs and other membrane proteins under the influence of various chemical and biological ligands is now an area that requires robust tools to establish quantitative trafficking phenotypes.

To tackle these challenges, fluorescent protein (FP) based sensors have been developed to study intracellular pH changes. (Super-)ecliptic pHluorin exhibits low fluorescence signal at pH < 6 and therefore has been widely used to investigate secretory vesicle exocytosis, internalization, and recycling.^9,10,11,12^ Ratiometric pHluorin produces a characteristic ratio with two excitation wavelengths and has been employed to measure the pH of various organelles.^13^ An enhanced, ratiometric pHluorin2^14^ was developed with increased fluorescence in acidic compartments. These characteristics made it a useful probe to study protein dynamics in endocytosis and recycling. While undergoing trafficking, the surface proteins are exposed to local environments with measurable pH changes that can be used to assess trafficking pathways. As an alternative to pH sensitive GFP, a Förster Resonant Energy Transfer (FRET)-based protein pH sensor family, pHlameleons^15^, were established using FRET from a pH insensitive CFP to various pH sensitive YFP variants. To further advance multicolor imaging capabilities, a red fluorescent protein pH sensor was developed and was shown to detect pH changes in the cytosol and mitochondria.^16^ Subsequently, pHuji^17^, an alternative red fluorescent protein pH sensor, with increased pH sensitivity, was demonstrated in tandem with pHluorin to image exocytosis and endocytosis of the transferrin receptor and B2AR simultaneously, demonstrating its utility as a second color pH probe.

A general weakness of FP based pH sensors, however, is that they label the total expressed pool of the protein of interest. After endocytosis the internalized pH sensitive FP-tagged receptors become indistinguishable from the resident intracellular pool of FP-tagged receptors, and contribute to background signal from incompletely quenched FPs or previously internalized protein in various acidic compartments. Discrimination of the surface pool from biosynthetic pools, which are also in neutral compartments, poses a fundamental limit for many FP pH sensor tags. Although FP methods have been developed to distinguish surface receptor localization vs. intracellular by using a surface FRET based assay^18^, this method only provides enhanced surface measurements without investigating individual vesicle trafficking after receptor activation.

Several non-FP methods have been developed to measure the pH of endosomal pathways and pH alterations in compartments. Programmable pH sensitive DNA nanomachines have been used as a tool to map endocytic pathways by tagging select trafficking proteins,^19^ and cysteine cathepsin selective cell-permeable bifunctional probes have been designed to detect specific combinations of pH and protein activity.^20^ However, for the purpose of selectively studying GPCR trafficking itineraries, using chemo-selective methods that selectively visualize cell surface receptors, rather than the active site binders or ligands that bind the receptor, is necessary to achieve high-contrast labeling of the plasma-membrane resident protein fraction, suitable for robust trafficking analysis under a variety of manipulations.

Current pH sensitive chemo-selective labeling methods either use expressed epitope tags for antibody binding, labeled-ligand binding, peptide tags that covalently link to modified fluorophores, or coiled-coil tag probes.^21,22^ However, these methods can involve lengthy incubation times for optimal labeling, and multiple wash steps to remove any unbound fluorophores or ligands, which can lead to higher background signal. Recent fluorogenic labeling applications reduced these challenges by using activatable “dark” fluorogens or fluorophores and show promising opportunities for superresolution microscopy.^23^ Development of far-red fluorogenic probes for superresolution imaging has improved the available photostability and brightness.^24,25^

We recently demonstrated rapid labeling of fluorogen activating protein (FAP) tagged B2AR at the cell surface using a cell-excluded fluorogenic ligand based on the far-red triarylmethane dye malachite green (MG-B-Tau).^26^ FAP technology utilizes a single chain antibody fragment that specifically binds and activates fluorogenic dyes that are otherwise non-fluorescent.^27^ This FAP-dye labeling approach allows for high specificity, rapid dye labeling, and does not require wash steps to remove unbound (dark) d ye. While the cell-excluded fluorogenic dye, MG-B-Tau, enabled detection of FAP-fused surface proteins, there remains a need for a robust method for dynamic imaging of receptor trafficking and pH changes during active endocytosis and recycling.

Here we demonstrate a family of far-red, genetically targetable, ratiometric, <u>a</u>ctivatable <u>pH</u> indicator <u>c</u>omplexes (TRApHIC), based on a pH sensitive Cy3 indicator dye linked to the pH independent fluorogen malachite green, bound to the dL5** FAP. In the absence of FAP binding, the MG moiety quenches emission from the cell-excluded tandem dye, but upon binding of MG to FAP the indicator complex is formed, displaying dual fluorescence excitation with far-red emission, and an excitation ratio that depends on the pH of the local environment. A family of pH sensitive Cy3 analogs provided a range of pKa values suitable for measurement of extracellular pH changes or changes in the endocytic/recycling pathway. The pH probe was validated in studies of endocytosis and recycling of a model GPCR, the β2-adrenergic receptor, in both confocal and Stimulated Emission Depletion (STED)-based superresolution imaging, confirming reported differences between agonists in induced endocytosis and revealing differences in recycling after agonist removal.

## RESULTS

### pH Sensor Development

Previously, a FAP-based TO1-cypHer5 pH sensor^28^ was established. When bound to the cognate FAP, the cell impermeable tandem dye based on thiazole orange and cypher5 displays pH dependent FRET and an increased ratio in the acceptor to donor emission signal. This sensor was used to study receptor trafficking endpoints and antigen transfer in dendritic cells. In this sensor system, the acceptor shows fluorescence signal at low pH in the absence of FAP, resulting in a modest, 10-fold fluorogenic activation upon binding and the potential for significant off-target fluorescence due to cellular uptake into acidic compartments. TO1-cypHer5 is an emission-based ratiometric sensor, requiring the separation of the two emission channels during image acquisition and careful evaluation of photobleaching of the independent channels to provide robust pH measurements. To overcome these limitations, we developed a new tandem dye sensor using malachite green (MG) as a fluorogenic acceptor, and a pH dependent Cy3-based donor. This sensor is a variant of our previously demonstrated Cy3-MG tandem dyes^29^ employed for signal enhancement, and establishes a genetically targetable, highly fluorogenic, excitation-based ratiometric pH reporter that enables quantitative measurement of physiologic pH changes during endocytosis and recycling, dynamically in living cells.

**Scheme 1.**
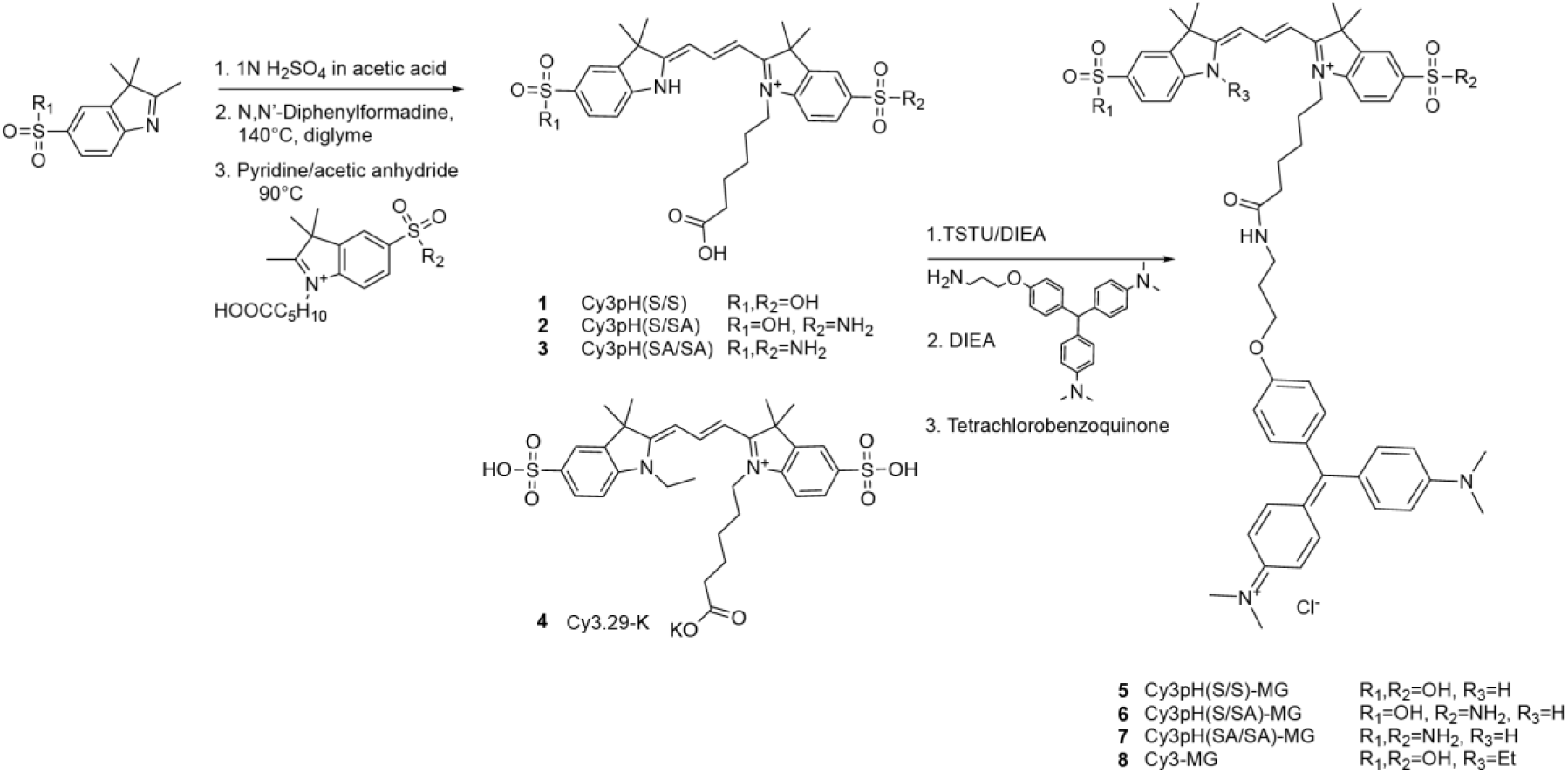
Synthesis of pH dependent cyanine-malachite green tandem dyes. A new series of unsymmetrical Cy3-based pH sensitive donors (**1-3**), prepared as active esters were coupled to an amine-modified MG analog to produce a series of pH sensor tandem dyes (**5-7**) (Scheme 1). In the same fashion, the pH-insensitive analog (**8**) was made from the potassium salt of Cy3.29, a bis-alkylated Cy3 analog (**4**). Upon binding of the MG fluorogen moiety to a FAP (dL5**), the quenched tandem dye becomes activated (Supplementary Fig. 1; Supplementary Fig. 2). In both free and FAP-bound states, the Cy3 signal is efficiently quenched, and the excitation is transferred to the free MG, which acts as a quencher, or to the MG-FAP fluorescent complex, which displays FRET-sensitized emission. At low pH, protonation of the indole nitrogen of the pH sensitive Cy3 moiety extends the resonant aromatic system, increasing the donor absorption and energy transfer signal. While detecting emission from the MG-FAP complex (680 nm), the excitation contribution of the pH sensitive moiety (**1-3**) increases monotonically with respect to pH, whereas MG direct excitation is relatively pH independent (illustrated in Fig. 1a and shown in Fig. 1b).

**Figure 1.**
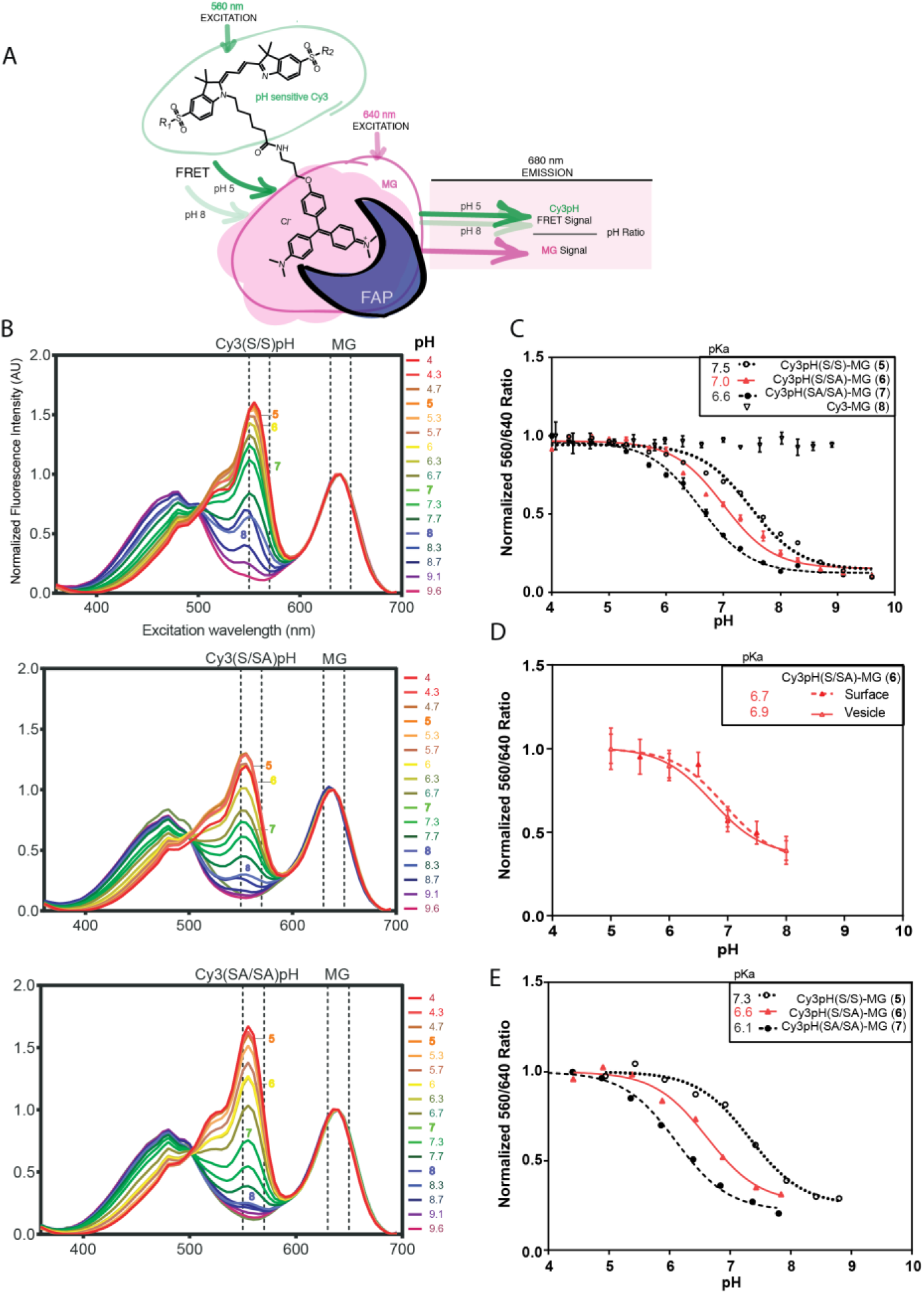
TRApHIC labeling results in pKa tunable, pH responsive ratiometric probes. A. Illustration of dye structure pH sensitivity and the excitation-based ratiometric pH response. B. Excitation spectra of Cy3(S/S)pH-MG, Cy3(S/SA)pH-MG, and Cy3(SA/SA)pH-MG from pH 4.0 to pH 9.6, complexed with dL5**. Emission was recorded at 710 nm in TECAN Infinite M1000 plate reader. C. Ratiometric characterization of dyes (**5**-**8**) response to pH. The normalized ratio of the fluorescence emission of the dye-protein complex in solution was measured using two excitation wavelengths (560 nm and 640 nm, 710 nm emission) was plotted against the solution pH. Mean results were fitted to a non-linear sigmoidal curve, error bars are shown as S.D. TRApHIC pKas were determined from curve fitting. D. Live FAP-B2AR expressing HEK cells were incubated with Cy3(S/SA)pH-MG and calibrated vs pH by nigericin/high K^+^ clamping. Data is shown as surface object measurements (no drug added) and vesicle object measurements (ISO induced endocytosis). The normalized ratio of the fluorescence emission from two excitation wavelengths (560 nm and 640 nm) was plotted against the solution pH. Mean results were fitted to a non-linear sigmoidal curve, error bars are shown as S.D. Dye pKas were determined from curve fitting. E. Live FAP-CFTR expressing HEK cells were incubated with each dye and the normalized ratio of the fluorescence emission from two excitation wavelengths (560 nm and 640 nm) was plotted against the solution pH. Results were fitted to a non-linear sigmoidal curve, pKas were determined from curve fitting.

The excitation ratio (560 nm/640 nm) of the family of dyes responds to pH changes with a characteristic sigmoidal curve, showing a ∼5-fold change in the ratio from pH 5 to pH 8 when complexed with purified dL5** protein in solution (Fig. 1c). The pK_a_ of the mono-sulfonamide analog (Cy3pH(S/SA)-MG (**6**), pKa = 7.0) is better suited for studying endocytosis and recycling than the bis-sulfonate analog (Cy3pH(S/S)-MG (**5**), pKa = 7.5), which shows less response throughout the physiological range and significant FRET at neutral pH. In addition, (**6**) is more useful than the bis-sulfonamide analog (Cy3pH(SA/SA)-MG (**7**), pKa = 6.6), showing the greatest response at moderately acidic conditions, typically sampled by recycling receptors. However, the bis-alkylated Cy3 donor (Cy3-MG, **8**), is unresponsive to pH changes.

All sensors tightly bind to the same FAP, providing a pH sensitive response that depends only on the dye. These dyes do not fluoresce in the absence of FAP and are cell impermeable (Supplementary Fig. 3; Supplementary Fig. 4). The availability of sensors with tunable pKa is useful for studying physiology in various contexts with a common genetically encoded tag. These probes bind to dL5** with high affinity (K_d_s < 10 nM), suitable for imaging on timescales from minutes to hours, and possess similar quantum yields (Table 1).

**Table 1.** Dye comparisons (**5**-**8**) with reported % quantum yield (QY), fold activation, pKa, and Kd.

| Dye | Quantum Yield (%) |  | Fold Activation |  | pKa | Kd (nM) |
| --- | --- | --- | --- | --- | --- | --- |
|  | pH = 5 | pH = 8 | pH = 5 | pH = 8 |  |  |
| Cy3(S/S)pH-MG (**5**) | 21.2 | 17.4 | 321 | 116 | 7.5 | < 10 |
| Cy3(S/SA)pH-MG (**6**) | 21.8 | 13 | 91 | 122 | 7 | < 10 |
| Cy3(SA/SA)pH-MG (**7**) | 21.2 | 10.4 | 67 | 49 | 6.6 | < 10 |
| Cy3-MG (**8**) | 25.1 | 16.2 | 136 | 110 | NA | < 10 |

### Cellular Calibration

To calibrate the sensor-FAP complex across physiological pH changes during receptor trafficking, the nigericin clamping method was used in live human embryonic kidney (HEK-293) cells stably expressing a dL5** FAP fused to the N-terminus of B2AR. The sensor (**6**) was first added to these cells to label B2AR on the plasma membrane through specific interaction with the extracellular dL5**. Cells were subsequently treated with 10 μM isoproterenol (ISO), a potent B2AR agonist, to induce endocytosis (vesicle measurements), or left untreated (surface measurements), followed by switching from the culture media to the nigericin calibration buffer with a high concentration of KCl. This treatment clamps the intracellular pH to the media pH, allowing ratiometric calibration of pH measurements in a living cellular milieu, whether on the surface or internalized into cellular compartments. The pH responsiveness of sensor (**6**) bound to dL5**-B2AR on the surface or vesicles analyzed by confocal microscopy both show the same sigmoidal titration curve as the in vitro analysis using dL5** complex with (**6**) in solution (Fig. 1d). Dyes (**5-7**) showed distinct vesicular calibration responses in living HEK293 cells expressing a dL5**-CFTR^30^ construct that paralleled the pKa values seen in the *in vitro* titrations, although shifted slightly lower in absolute fit pKa (Fig. 1e). The pH titrations of the proteins in cells hence parallel the response seen free in solution, demonstrating the tuning and cellular compatibility of this family of pH sensor dyes.

We evaluated a variety of imaging protocols to assess photobleaching during the imaging time course, recognizing there is a tradeoff between exposure time, intracellular dynamics, experimental duration, and ratiometric quantitation. After (**6**) surface labeling, B2AR endocytosis was induced with 10 μM isoproterenol (ISO) for 15 min, and then nigericin pH clamped in different pH buffers. Cells were imaged with exposure times of 100 ms or 500 ms (Supplementary Fig. 5) for 60 frames. Objects ≤1 μM diameter were identified, and quantified for excitation ratio (560 nm/640 nm) with a common emission filter and camera settings. 60 imaging cycles with 500 ms exposure for each channel (60s total illumination) resulted in a 10% change only at pH 5, potentially a result of the increased extinction coefficient of the donor at low pH coupled with the longer exposure. To eliminate potential photobleaching artifacts, using a short exposure time is preferred in live, real-time imaging ≥ 60 frames. Although the endocytosed vesicles are relatively immobile in the nigericin photobleaching experiments and endpoint experiment imaging, vesicles are more mobile during real-time receptor endocytosis and recycling, which may result in motion artifacts upon longer exposures that skews the ratio analysis without further image analysis and correction. In super resolution STED imaging with live cells (*vide infra*), endpoint imaging is optimal for reducing the total of repeated high intensity laser exposure, vesicle motion, and focal drift.

### B2AR Agonist-Induced Trafficking

Upon addition of the sensor dye, cells showed strong fluorescence staining on the plasma membrane in the 640 nm excitation channel. The surface 560/640 ratio reported the pH as 7-8, showing pH 7.5 as the average of the fitted Gaussian peak of object ratios (Fig. 2c), consistent with the pH of the imaging media. Soon after exposure to the agonists isoproterenol (ISO) or epinephrine (EPI), endocytic vesicles started to emerge from the cell surface and move inward (Supplementary Video 1, 2; Fig. 2 a,b ISO and EPI). The intensity of the plasma membrane decreased significantly and punctate vesicular structures gradually accumulated in the cytoplasm. Fluorescence intensities of the vesicles rose rapidly in both channels, consistent with an enrichment of the receptors in the vesicle following receptor clustering and fusion of endocytosed vesicles during vesicle trafficking. In addition, observable pH changes accompanied the redistribution of the receptors following endocytosis. ∼15 minutes into endocytosis, the majority of plasma membrane signal had dropped below the detection threshold. At this point, most of the detected objects were receptors in vesicular compartments. After 30.5 minutes, the pH of internalized vesicles covered a broad range, but had significantly decreased to an average of pH 6 for both ISO and EPI conditions (Fig. 2c ISO and EPI). This pH drop suggests continuous acidification of the population of vesicles in the sustained presence of an agonist. Due to the 30 second inter-frame interval, it was not possible to track single vesicles to assess temporal changes in pH, although such studies may be possible when a narrower time window is utilized for imaging. NorEPI displayed a different endocytic behavior than ISO and EPI, with a widespread population of receptors on the plasma membrane and a lesser amount in endocytosed vesicles by time point 30.5 min. (Supplementary Video 3; Fig. 2 a,b,c NorEPI). Over the complete time course, ISO and EPI had a higher population % in pH ≤ 6 than NorEPI treated cells and the negative controls (Figure 2b). Overall, the calculated t_1/2_ median endocytotic rate was ISO 5.76 (+/- 0.10) min, statistically indistinguishable from EPI at 5.58 (+/- 0.08) min, and lastly NorEPI at 11.8 (+/- 0.25) min (Table 2). The negative controls showed no signs of endocytosis with ≥ 80% objects at pH 7 and ≥ 12% at pH 8, and no significant shift from initiation to 30.5 min (Supplementary Video 4, 5; Fig. 2a,b,c ALP and DMSO).

**Figure 2.**
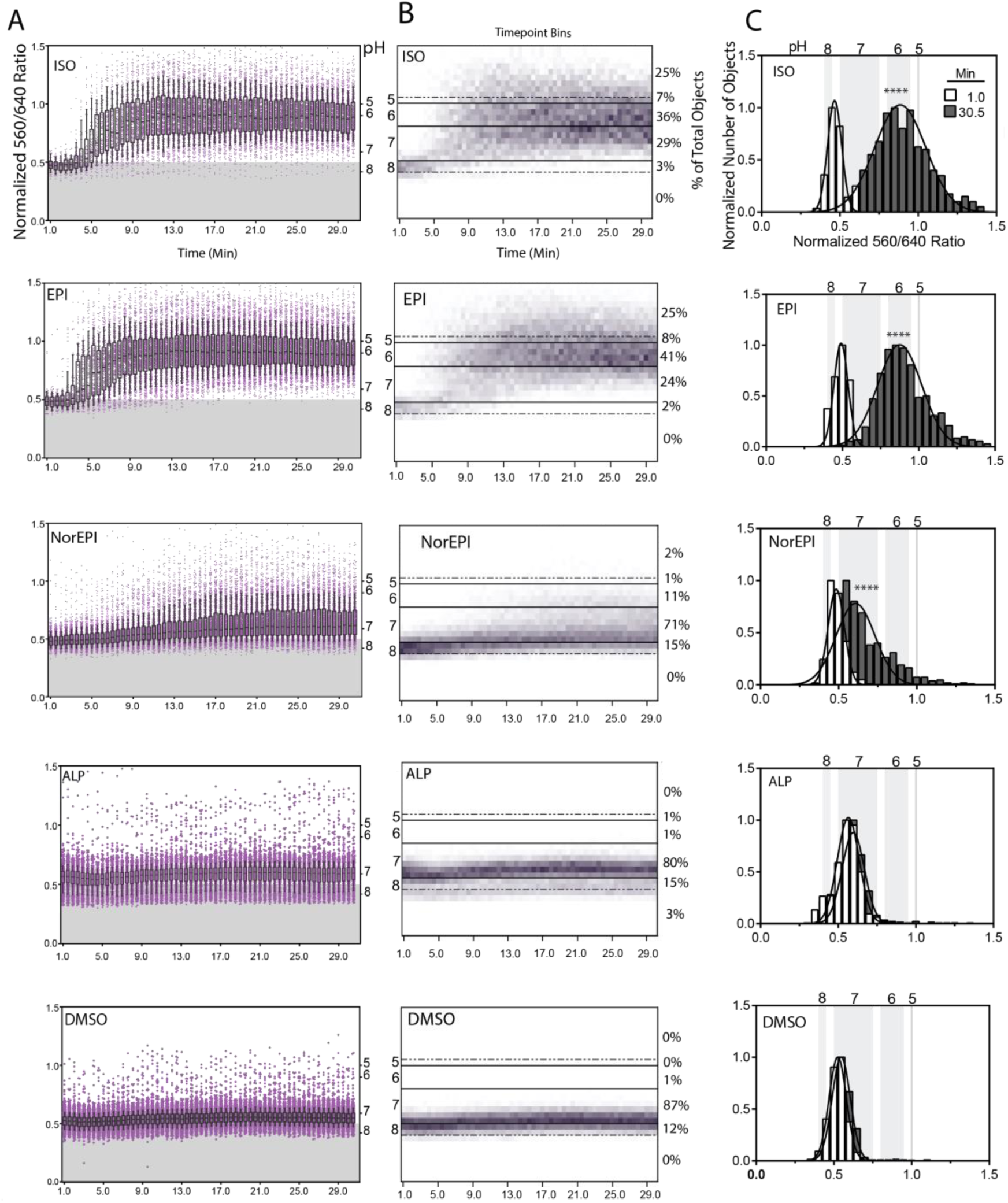
TRApHIC labeling highlights differences among B2AR agonists in endocytosis rates by single vesicle measurements. A. Analysis of individual vesicles from dL5**-tagged B2AR in real time. Each point represents a single vesicle. The normalized ratio of the emission fluorescence from the two-excitation wavelengths (560 nm and 640 nm) of single vesicles is shown on the left Y-axis and the pH from the single vesicle calibration on the right Y-axis. For the endocytosis experiment, either cells were dosed with 10 μM ISO, 10 μM EPI, 300 μM NorEPI, 10 μM ALP, or 1:1000 DMSO. 1 minute after addition, endocytosis was observed for 30.5 min with 30 sec intervals. Data includes boxplots representing 90^th^, 75^th^, median, 25^th^, and 10^th^ percentile. B. Heat-map and time point bins of vesicle results in A. pH bins are listed on the left Y-axis. Over the whole imaging time period, the total % objects at different pHs are listed on the right Y-axis. C. Histograms of time point 1 min and time 30.5 min vesicle ratios, curves are Gaussian fitted. The average Gaussian fit of each time point was compared. (T-test two tailed) ****P ≤ 0.0001; ***P≤ 0.001; **P≤ 0.01;*P≤ 0.05.No indicator P-value > 0.05

Recycling was observed after washing out agonist and adding antagonist, Alprenolol (ALP) (10 μM), which halted residual endocytosis. At the beginning of exocytosis, ISO was visible as 2 distinct populations, with an average of pH 6.6 and pH 5 (Supplementary Video 6; Fig. 3c ISO). EPI had a broad spread with an average of pH 6 (Supplementary Video 7; Fig. 3c EPI). During the recycling phase, the pH shifted back towards baseline pH (Fig. 3a,b,c ISO and EPI). As the vesicles moved back and fused with the plasma membrane, the plasma membrane regained its fluorescence signal and signal ratio associated with neutral pH. However, EPI showed 2 populations, one remaining with an average of pH 6, and the other restored to pH 7 (Fig. 3c EPI). NorEPI exhibited markedly different exocytosis behavior (Supplementary Video 8). At the start of NorEPI exocytosis, a large population of objects was detected in the pH 7 range (a sum of 2 Gaussian fitted curve with an average of 7.3 and 6.6) (Fig. 3c NorEPI). Over the exocytosis time course, the more acidic population quickly returned to neutral pH 7.2.(Fig. 3b,c). The t_1/2_ average exocytosis rates are shown in Table 2. Throughout the exocytosis imaging period, DMSO and ALP negative controls showed no significant change after washout and addition of ALP (Supplementary Video 9,10; Fig. 3a,b,c). The NorEPI treated cells showed significantly faster exocytosis and slower endocytosis than other agonists, suggesting that the agonist effect of the NorEPI may result in distinct signaling pathways that regulate internalization, intracellular signaling, and recycling.^31^

**Figure 3.**
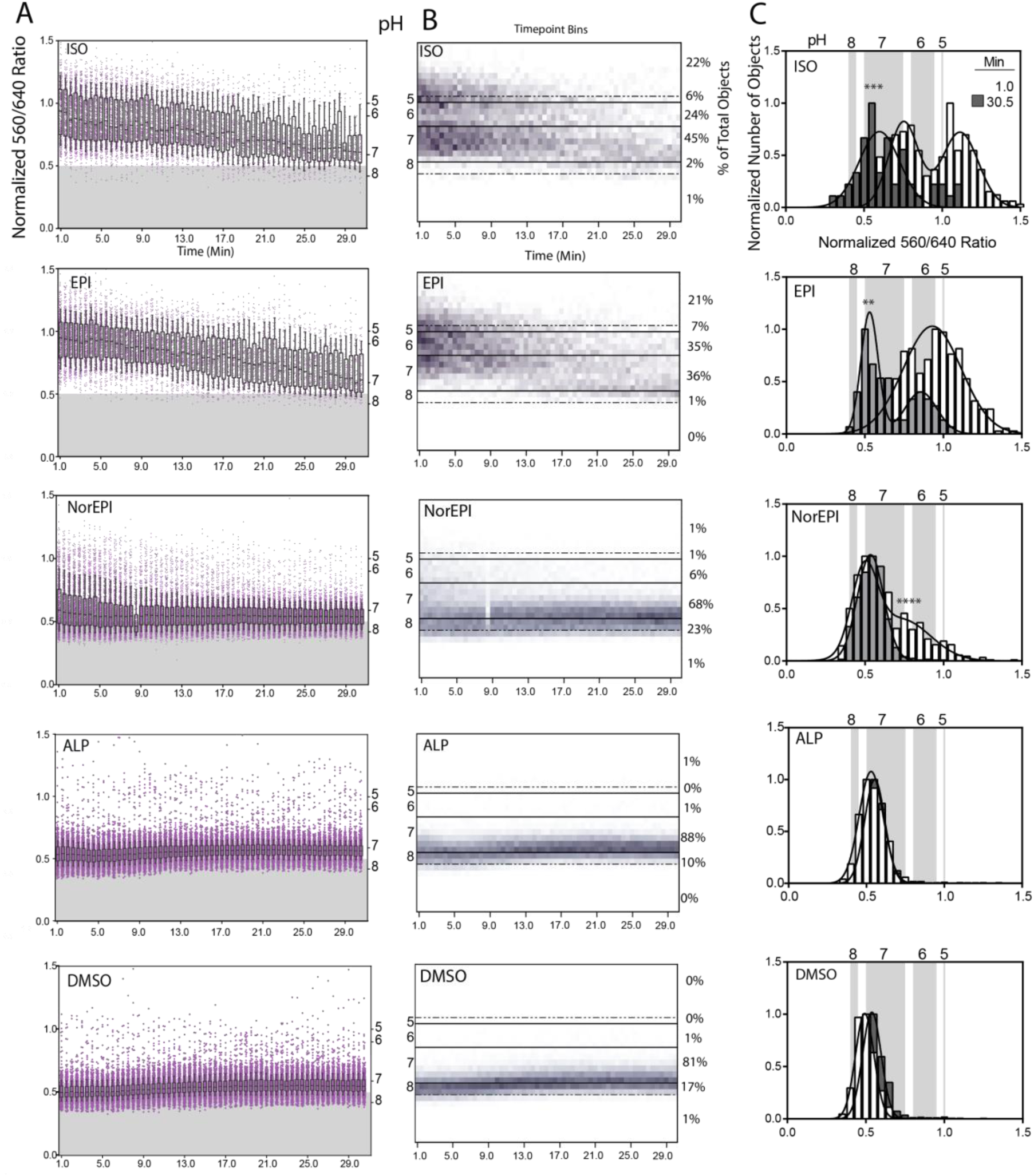
TRApHIC labeling highlights differences among B2AR agonists in recycling rates after drug removal and antagonist treament by single vesicle measurements. A. Analysis of individual vesicles from dL5**-tagged B2AR in real time. Each point represents a single vesicle. The normalized ratio of the emission fluorescence from the two-excitation wavelengths (560 nm and 640 nm) of single vesicles is shown on the left Y-axis and the pH from the single vesicle calibration on the right Y-axis. After endocytosis imaging, drug was washed out and 10 μM ALP was added to the cells and time lapse imaging started again to capture the receptor recycling process. Images were acquired every 30 sec for a duration of 30 min Data includes boxplots representing 90^th^, 75^th^, median, 25^th^, and 10^th^ percentile. B. Heatmap and time point bins of vesicle results in A. pH bins are listed on the left Y-axis. Over the whole imaging time period, the total % objects at different pHs are listed on the right Y-axis. C. Histograms of time point 1 min and time 30.5 min results, curves are Gaussian fitted. The average Gaussian fit of each time point was compared. (T-test two tailed). For results with 2 different populations for a timepoint, One-Way ANOVA multiple comparisons test was performed. ****P ≤ 0.0001; ***P≤ 0.001; **P≤ 0.01;*P≤ 0.05.No indicator P-value > 0.05

### Superresolution Ratiometric Physiological Imaging

Because the emission of the tandem dye occurs through the MG fluorogen regardless of excitation, we thought it probable that STED could be used to achieve superresolution pH measurements using dual excitation and a single depletion wavelength (775 nm), suitable for depletion of the MG fluorogen (Supplementary Fig. 6). Exclusively using the 775 nm depletion laser is ideal for reducing cellular damage, autofluorescence, light scattering, and photobleaching.^25^ We used a commercially available Leica STED system with dual excitation and 775 nm depletion for both excitation channels, and examined the resolution enhancement and ratiometric properties of FAP-B2AR labeled with tandem dye (**6**) in living cells. Figure 4a shows a confocal vs STED single frame image of endocytosed FAP-B2AR 20 min after dye labeling and ISO addition. The confocal and STED images show clear resolution differences that are more pronounced in select endosomal line plots (Fig. 4b). The line plots of the 560 and 640 ex channels in the confocal and STED channels show different ratiometric intensities, attributed to the differences in laser power between microscopy methods, required for optimal image quality. Simple raw confocal and STED object 560/640 ratios were obtained and plotted against each other, showing a positive linear correlation (R^2^ = .6), suggesting a robust ratiometric pH sensor response for both confocal and STED modalities. (Fig. 4c)

**Figure 4.**
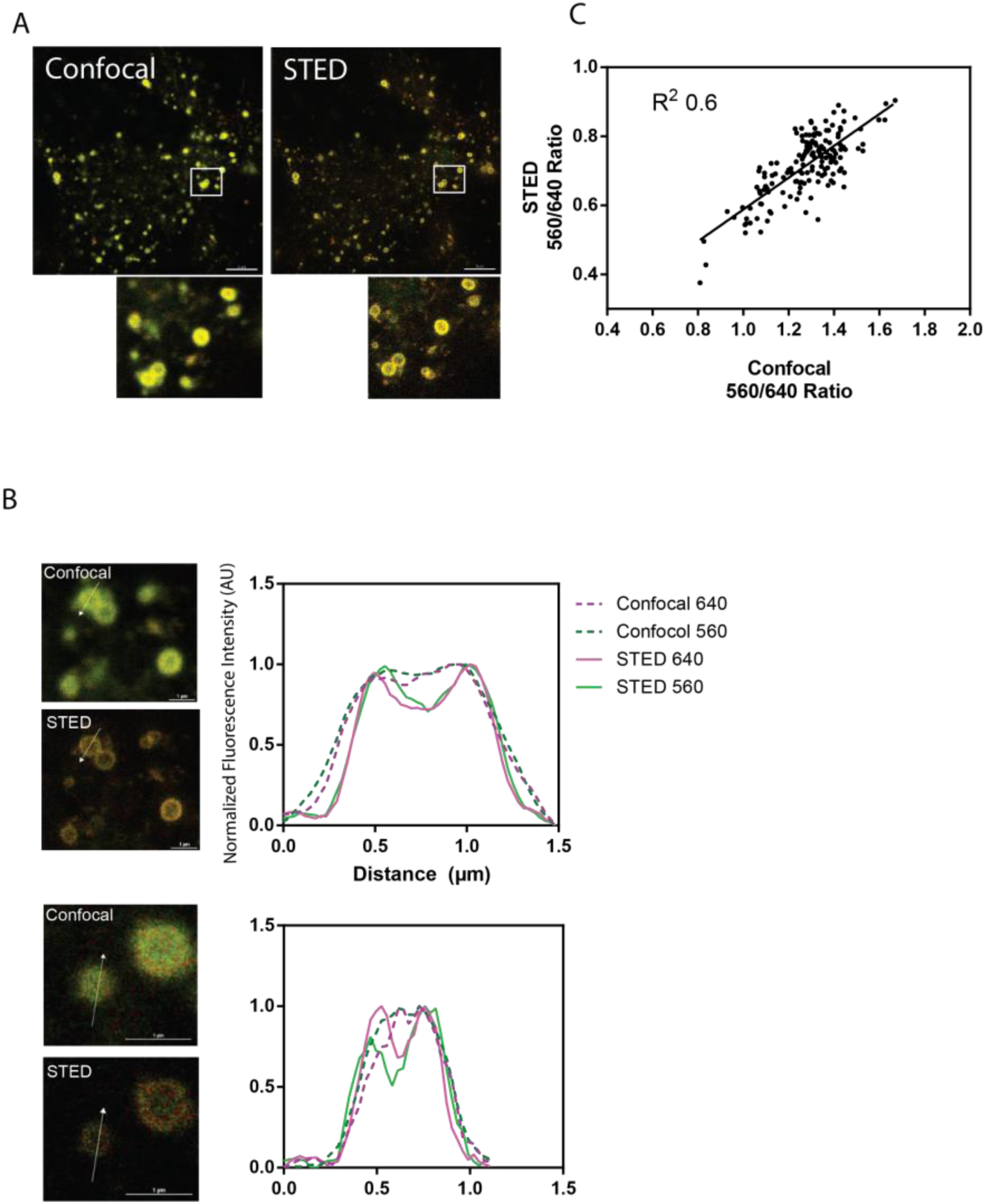
TRApHIC labeling is compatible with ratiometric STED-based superresolution imaging. A. Confocal and STED image of Cy3(S/SA)pH-MG labeled FAP-B2AR. Scale bar 5 μm. B. Line plots across individual endosomal objects in 560 and 640 excitation channels, show pronounced resolution enhancement under STED excitation. Scale bar 1 μm. C. STED endosomal object 560/640 ratio plotted against the confocal 560/640 ratio. The resulting ratios show a (R^2^ = .6) positive linear correlation with each other. (objects were collected from 3 different fields of view)

## DISCUSSION

We devised a new approach to develop highly fluorogenic tandem dye sensors, and demonstrated the utility of the approach using a series of tuned pH sensors to visualize for receptor trafficking. These sensor dyes are activated on binding with high affinity to a cognate FAP, genetically fused to a target protein, generating FRET based excitation ratiometric properties. These pH sensors are well-suited for monitoring the pH dynamics associated with endocytosis and recycling of receptors or other cell surface proteins. Surface exposed receptor labeling is achieved by adding the dye to the cells in media without washing, where the cell-impermeant dye selectively labels proteins at the plasma membrane. We demonstrated this labeling approach using agonist-mediated internalization of B2AR, providing a new approach to studying real time trafficking of membrane proteins in living cells and demonstrating the first excitation-ratiometric physiological indicator useful in STED microscopy.

These probes are well-suited for measuring a range of pH changes in receptor trafficking, or measuring pH changes in the extracellular and pericellular space. The dyes used as physiological indicators in these sensors are chromogenic and fluorescent in low pH environments, resulting in acquisition of a new excitation channel as a result of physiological changes. It is possible that other reversible or reaction-based indicators could also be paired with a quenching fluorogen acceptor to produce new targeted, ratiometric, activatable physiological sensors for use in living cells. Although the pH sensitive probes demonstrated here are cell-excluded, other sensor dyes could yield cell-permeant physiological sensors, although the overall molecular weight may limit the ability of some sensor dyes to access intracellular targets.

## METHODS

### Plasmid and Cell Line Generation

pBabe-dL5**-β2-AR was generated previously^24^ and plasmid is available at Addgene (ID 101253). Stable HEK293 cells were generated by transfecting HEK293 cells with pBabe-dL5**-β2-AR followed by drug selection (1 mg ml^−1^puromycin, Invitrogen) and FACS enrichment (Becton Dickinson FACS Vantage flow cytometer. Excitation: 633 nm; emission: 685/35 nm).

### Cell Culture

HEK cells were cultured in Dulbecco’s Modified Eagle Medium. Cultures were supplied with 10% fetal bovine serum and incubated at 37 °C in 5% CO_2_.

### pH Calibration Buffer for In-vitro Measurements

In order to keep ion composition of the pH buffers the same across the whole range of the pH that should be covered several weak acids were used. 4.8g of glacial acetic acid, 7.84 g of phosphoric acid and 4.95g of boric acid were dissolved in 150mL of DI water and brought up to 200 mL total making the stock acidic solution. For each pH buffer 12.5 mL of the acidic stock was transferred to 50 mL tube titrated with 2M NaOH to the desired pH and brought to 50 mL total.

### Spectroscopic Analysis

UV-Vis absorption: roughly 0.1 mg Cy3pH(S/SA)-MG was dissolved in ethanol with 1% acetic acid to prevent carbinol formation of MG. The tandem dye concentration was measured by the absorbance peak of MG at 606 nm (ε = 91,600 M^−1^cm^−1^) in acidic ethanol using a UV-Vis spectrophotometer (PerkinElmer).

### Excitation Spectra

Dyes with 5-fold excess of dL5 were precomplexed in pH buffers from pH 5 to pH 9 keeping sample absorption below 0.4 at main absorption peaks. Excitation scan was performed from 400 nm to 680 nm with the emission recorded at 710 nm and measured on a TECAN Infinite M1000 fluorescence plate reader.

### Fluorescence Titration

Secretion and purification of soluble dL5 was as described previously.^32^ Binding affinity to soluble dL5 was measured on a TECAN Infinite M1000 fluorescence plate reader. 1 nM dL5 was mixed with varying concentrations of dye at neutral pH (7.4) in PBS. The FAP+dye complex fluorescence was corrected by subtracting the fluorescence of a dye only sample. The data was fitted using a binary ligand depletion equilibrium model in Prism 6.0, which reported kD. Data points are the mean ± S.D. from triplicate measurements.

### Live Cell Spinning Disk Confocal Microscopy

500 nM Cy3pH(S/SA)-MG (**6**) was added to cells to label dL5**-B2AR for 15 min. Then either 10 μM isoproterenol (ISO) (Cayman Chemicals), 10 μM epinephrine (EPI) (Cayman Chemicals), 300 μM norepinephrine (NorEPI) (Cayman Chemicals), 10 μM alprenolol (ALP) (Cayman Chemicals), or vehicle DMSO, was added to the cells and time lapse imaging started 1 minute after the addition of drug. Images were acquired every 30 sec for a duration of 30.5 min. Then drug was washed off by rinsing cells with at least 4 volumes of imaging medium. After washout, 10 μM ALP was added to the cells and time lapse imaging started again to capture the recycling process. Images were acquired every 30 sec for a duration of 30.5 min. During imaging, cells were cultured in Opti-MEM (Invitrogen) in Mattek dishes (MatTek Corp.) and kept in an imaging chamber at 37 °C with 5% CO2 supply (Pathology devices). Cells were imaged on an Andor Revolution XD system (Andor technology) with a Yokogawa CSU-X1 spinning disk confocal unit (Yokogawa Industries). Cells were imaged with a 60x, 1.49 TIRF objective (Nikon) and excited with a 640 nm and a 560 nm laser sequentially. Emission was collected with a 685/70 band pass for both channels. EM gain was set at 300.

### Live Cell Leica STED and Confocal Microscopy

500 nM Cy3pH(S/SA)-MG was added to cells to label dL5**-B2-AR for 15 min. Then 10 μM isoproterenol (ISO) (Cayman Chemicals) was added to the cells and imaging started 20 min after addition. Imaging was done on Leica microscope with 100x/1.40 HC PL APO oil objective. Cells were excited with 561 nm and 633 nm laser excitation using a white-light laser and the acousto-optic beam splitter respectively, while emission was detected at 660 nm - 725 nm (gain 100, time gate of 0.5 ns-6.5 ns). STED imaging utilized 775 nm depletion laser (power 100%) for both excitation channels. Confocal laser power: 561ex (56%), 633 ex (12%); STED laser power: 561 ex (100%), 633 ex (64%). During imaging, cells were cultured in Opti-MEM (Invitrogen) in Mattek dishes (MatTek Corp.) and kept in an imaging chamber at 37 °C with 5% CO2 supply.

### pH Calibration with Nigericin Clamping

Nigericin calibration buffers contained 140 mM KCl, 5 mM α-D-glucose, 0.5 mM CaCl_2_•2H_2_O, 1 mM MgCl_2_ and the pH (pH 5 to pH 8) of the mixture was adjusted using 20 mM MES and 20 mM Tris base, where MES and Tris base were added in different ratios to generate different pH buffers.^33^ Cells were treated with the calibration buffer with 10 μM Nigericin (Sigma) for 10 min prior to imaging.

### Single Object Ratio Analysis

Analysis was carried out using Imaris software. Objects were identified with spot detection and background subtraction in the 640 nm channel. After identifying objects, they were automatically thresholded based on the “average center intensity” in the 640 nm channel using default settings. The resulting object’s average intensity measurements were determined in the Cy3pH(S/SA) FRET (560 nm ex) and MG (640 nm ex) channels. For each identified object, the average 560/640 ratio was plotted vs. time after treatment for endocytosis and recycling conditions (Fig. 2a; Fig. 3a). The 75^th^, median, and 25^th^ percentile were calculated at every time point and median was fitted to a sigmoidal curve to determine the t_1/2_ values (curve not shown, all curves R^2^>.85). The boxplot whiskers represent the 90^th^ and 10^th^ percentile. The experimental data was also binned to .05 intervals from 0 −1.5 (normalized 560/640 ratio) for each time point. Collectively, a heat map was generated, which displays the consensus and what percentage of all identified objects are in a pH range (Fig. 2b; Fig. 3b). The first time point (1 min) and last time point (30.5 min) bins for both endo- and exocytosis experiments, were plotted as a histogram and fitted to a robust Gaussian, or sum of 2 Gaussians, curve (Fig. 2c; Fig. 3c). The histogram fitted curves demonstrate if there was a significant population shift in pH between selected time points. Data analysis was performed with Prism 6.0. ****P ≤ 0.0001; ***P≤ 0.001; **P≤ 0.01;*P≤ 0.05.No indicator P-value > 0.05

### pH Calibration Analysis

Expressing cells were labeled with 500 nM Cy3pH(S/SA)-MG for 15 min and then subjected to either 10 μM ISO (vesicle) for 15 min or no drug (surface). In both conditions, nigericin clamping was done at each pH (from pH 5 to pH 8), at least 6 pairs of images (640 nm excitation followed by 560 nm excitation, both 680 emission) were taken for the ratio analysis. Exposure time was 100 ms for both channels. Objects were combined from all the images and binned histograms representing the distribution of the ratio were generated by single Gaussian peak fitting. The best-fit average values were used for plots and non-linear sigmoid fitting using Prism 6.0. Data points are the mean ± S.D. from the fit.

### Photostability Measurement

Cells were labeled with 300 nM Cy3pH(S/SA)-MG and exposed to 10 μM ISO for 15 min. Then the cells were subjected to nigericin clamping. Following Nigericin clamping, the cells were imaged with 640 nm excitation and 560 nm excitation sequentially for 60 cycles with an exposure time of 100 ms or 500 ms for both channels. The data is expressed as the mean ± S.E.M. of 3 different fields of view.

### Statistical Analysis

All statistics were performed by GraphPad Prism 6.0 and are either described in methods or in figure legends.

### Data Availability

Synthesis and NMR of the final products is available in the supporting information. Cy3(S/SA)pH-MG cell impermeability and specificity is shown in supporting information. All data presented are available from the corresponding author upon request.

## FUNDING RESOURCES

NMR instrumentation at CMU was partially supported by NSF (CHE-0130903 and CHE-1039870). The STED microscope was acquired under the NIH shared instrument grant program S10OD021540. This work was supported in part by NIH grants R01EB017268 and U54GM103529.

## ACKNOWELDGEMENTS

We thank Dr. Manojkumar Puthenveedu and Zara Y. Weinberg for advice with B2AR trafficking, Ulf Schwarz for assistance with STED microscopy, and Dr. Cheryl Telmer for helpful discussion.

## AUTHOR INFORMATION

Author Contributions

L.A.P performed imaging experiments, data analysis, and wrote the paper. Q.Y. performed imaging, photobleaching experiments, initial data analysis, and wrote the paper. B.F.S synthesized TO1-Cypher5, Cy3-MG, Cy3pH(S/S)-MG, Cy3pH(S/SA)-MG, and Cy3pH(SA/SA)-MG. D.K. developed quick pipeline for organizing large single-vesicle time-course data sets for Prism, performed spectroscopy data collection, and helped with methods. M.B.L. performed pH calibration and data analysis. S.C.W. did data analysis and provided assistance with STED microscopy. L.K. and S.S. performed emission and excitation spectra of Cy3pH(S/S)-MG, Cy3pH(S/SA)-MG, and Cy3pH(SA/SA)-MG. M.P.B. designed sensors, experiments, and wrote the paper.

## COMPETING INTERESTS

M.P.B. is founder and Chief Scientific Officer at Sharp Edge Labs, Inc., a licensee commercially utilizing the FAP-fluorogen technology.

